# An alveolus lung-on-a-chip model of *Mycobacterium fortuitum* lung infection

**DOI:** 10.1101/2024.08.30.610530

**Authors:** Victoria Ektnitphong, Beatriz R.S. Dias, Priscila C. Campos, Michael U. Shiloh

## Abstract

Lung disease due to non-tuberculous mycobacteria (NTM) is rising in incidence. While both two dimensional cell culture and animal models exist for NTM infections, a major knowledge gap is the early responses of human alveolar and innate immune cells to NTM within the human alveolar microenvironment. Here we describe development of a humanized, three-dimensional, alveolus lung-on-a-chip (ALoC) model of *Mycobacterium fortuitum* lung infection that incorporates only primary human cells such as pulmonary vascular endothelial cells in a vascular channel, and type I and II alveolar cells and monocyte-derived macrophages in an alveolar channel along an air-liquid interface. *M. fortuitum* introduced into the alveolar channel primarily infected macrophages, with rare bacteria inside alveolar cells. Bulk-RNA sequencing of infected chips revealed marked upregulation of transcripts for cytokines, chemokines and secreted protease inhibitors (SERPINs). Our results demonstrate how a humanized ALoC system can identify critical early immune and epithelial responses to *M. fortuitum* infection. We envision potential application of the ALoC to other NTM and for studies of new antibiotics.

## Introduction

Nontuberculous mycobacteria (NTM) represent a diverse group of environmental organisms found in varied natural habitats such as soil, water, and vegetation (Mercaldo et al., 2023). Traditionally considered opportunistic pathogens of people with structural or immune deficits, lung disease secondary to NTM is increasing in incidence in women and people over 65 years of age (Winthrop et al., 2020). Among the NTM, *M. fortuitum* is the second most frequently encountered rapidly-growing NTM after *Mycobacterium abscessus*, and can cause a spectrum of infections including pulmonary, skin and soft tissue, and disseminated infections (Roquet-Baneres et al., 2023, Kim et al., 2023). NTM lung infections are more frequent and more severe in immunocompromised individuals and individuals with pre-existing conditions such as bronchiectasis, severe chronic obstructive pulmonary disease (COPD), cystic fibrosis, organ transplant and cancer (Roquet-Baneres et al., 2023, Kim et al., 2023). Treatment of NTM lung disease can be complex due to the presence of antibiotic resistance, medication side effects and extended duration of drug treatment. More efforts are needed to develop new, safe and efficacious therapeutics and to shorten treatment duration (Johansen and Kremer, 2020, Orme and Ordway, 2014, Kremer, 2020). In addition, the rising incidence of NTM lung infections among individuals without defined predisposing conditions indicates that a greater understanding of the immune response to NTM lung infection is needed.

Expanding the scope of knowledge of the dynamic interactions between humans and NTM is essential for developing effective prevention and treatment strategies for NTM lung disease, but gaining such knowledge has been hindered by the lack of suitable *in vitro* and *in vivo* models of NTM lung infection (Baldwin et al., 2019). Current *in vitro* models typically use traditional two-dimensional cell culture methods with primary human CD14^+^ peripheral blood-derived macrophages or immortalized monocytic cell lines THP-1 and U937 (Kilinc et al., 2022), and alveolar epithelial cell lines such as A549 (Rampacci et al., 2020). Although immortalized cell lines allow for large-scale screening and assay reproducibility, a major disadvantage is that they may not completely replicate myeloid cell responses in humans, especially in the unique alveolar microenvironment. For example, the use of PMA to differentiate THP-1 and U937 cells into adherent macrophage-like cells alters monocytic surface markers, transcriptional activities, and cytokine production (Rampacci et al., 2020). One recent advance beyond traditional two-dimensional systems has been adaptation of primary human airway-derived organoids to the study of *M. tuberculosis* and *M. abscessus* pathogenesis and drug susceptibility (Alcaraz et al., 2022, Iakobachvili et al., 2022, Leon-Icaza et al., 2023, Kim et al., 2024). Animal models have also been established to study host immune responses to NTM and for testing potential antimicrobial compounds and vaccines. Most animal models of NTM lung infection use mice as the host species including for *M. abscessus* (Caverly et al., 2015, De Groote et al., 2014) and *Mycobacterium avium* (Gonzalez-Perez et al., 2013, Gangadharam, 1995). However, the variable virulence of NTM in model organisms makes establishing consistent animal models challenging. For example, *M. avium* can establish a productive lung infection in immunocompetent mice, whereas *M. abscessus* is typically rapidly cleared and requires immunocompromised mice to achieve a similar outcome (Chan et al., 2016). Animal models for other NTM species such as *M. fortuitum* are lacking, delaying progress towards understanding shared and unique molecular mechanisms of pathogenicity for various NTM. Thus, developing *in vitro* models that can recapitulate the human lung microenvironment using primary human cells has the potential to advance the study of pulmonary NTM infections, and is a critical step towards developing improved prevention, vaccine and treatment strategies.

The alveolus lung-on-a-chip (ALoC) is an innovative microfluid device composed of microfabricated channels and chambers that mimic the architecture and cellular composition of the lung alveolus, the initial site of interaction for pulmonary NTM infection (Bethencourt et al., 2024). The “chip” contains two overlapping channels, parallel to each other, that are separated by a porous membrane. It features alveolar epithelial cells lining one channel and pulmonary microvascular endothelial cells (HMVECs) lining the other to closely replicate the alveolar-capillary interface of the human lung (Jain et al., 2018). Because the cell types grow in distinct channels, media or air flow through the channels can be independently controlled to create distinct environments for the cells. Additional advantages of the ALoC include the ability to generate an air-liquid interphase (ALI) that is hallmark of the alveolus and application of mechanical stretch that is essential for the proper differentiation of alveolar epithelial cells into type I (AT1) and type II (AT2) pneumocytes. In that respect, the ALoC helps overcome one of the major obstacles of *in vitro* studies of alveolar epithelial cells, namely, simultaneously differentiating both AT1 and AT2 cells in culture by leveraging stretch and growth factor addition (Li et al., 2018). Thus, compared to traditional tissue culture methods, the ALoC system provides a physiologically relevant environment to study cells and pathogens of the human lung and has been applied to several pathogens including *E. coli* (Huh et al., 2010), *M. tuberculosis* (Thacker et al., 2020), *Aspergillus fumigatus* (Hoang et al., 2022), *Staphylococcus aureus* (Bai et al., 2022, Deinhardt-Emmer et al., 2020), and influenza virus (Bai et al., 2022, Deinhardt-Emmer et al., 2020).

The complex interaction between *M. fortuitum* and the human alveolar epithelial and immune cells remains poorly understood. Here, we develop and use a fully humanized ALoC model as a platform to study the host-pathogen interactions of *M. fortuitum* within the alveolar microenvironment. By recapitulating key features of the alveolar interface, including cellular composition, mechanical forces, and biochemical gradients, this model offers a unique opportunity to investigate the dynamics of *M. fortuitum* infection in a controlled and reproducible manner. Through integration of advanced imaging techniques and transcriptomic analyses, we identify macrophages as the primary cell type infected in the alveolus and upregulation of several inflammatory pathways after *M. fortuitum* infection of the humanized ALoC. We propose application of the humanized alveolus lung-on-a-chip as a new model to study NTM infections.

## Results

Two types of lung chips have been developed to date, one that approximates a bronchial airway (Benam et al., 2015, Benam et al., 2016) and another more like an alveolus (**Figure 1A**) (Huh et al., 2012a, Huh et al., 2010, Huh et al., 2012b). Since early NTM infection occurs in the alveolus, we chose the alveolus lung-on-a-chip (ALoC) as our working model for NTM infection (**Figure 1B**), as previously demonstrated with *E. coli* (Huh et al., 2010) and *M. tuberculosis* (Mishra et al., 2023, Thacker et al., 2020). We initially decided to use *M. fortuitum* to model airway NTM infection using the ALoC model both because it is rapidly-growing, with a doubling time of 2-3 hours (Qin et al., 1999), and because amongst the rapid growers it has one of the highest age-adjusted incidence of disease (Donohue, 2021).

**Figure 1.**
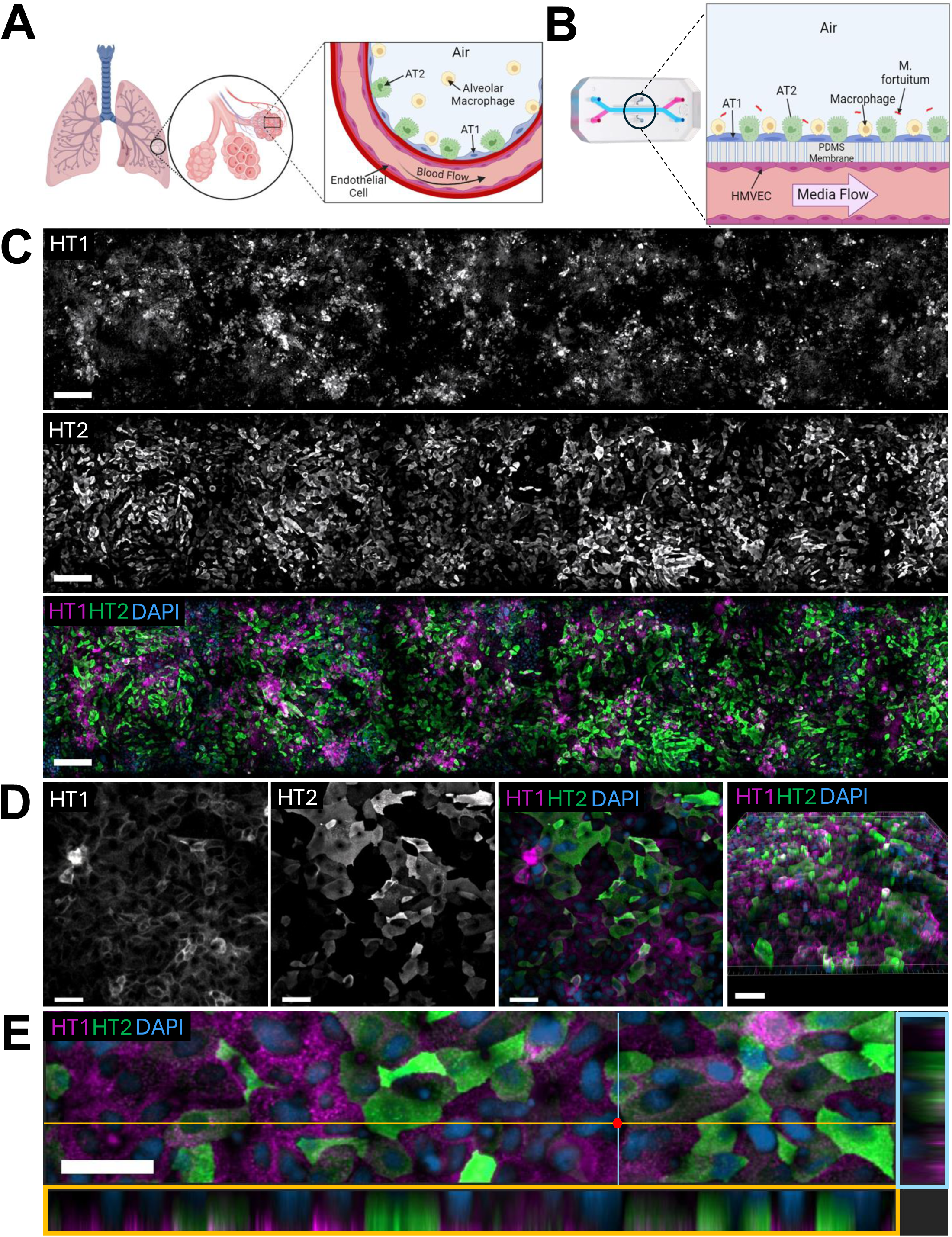
Establishment of a fully humanized alveolus lung-on-a-chip (ALoC) model system. **A.** Schematic of a human alveolus and the surrounding vasculature, highlighting alveolar epithelial type I (ATI), alveolar epithelial type II (AT2) cells, and alveolar macrophages in the alveolar sac, with pulmonary microvascular endothelial cells lining the vasculature. **B.** Schematic of the ALoC model for *M. fortuitum* infection. The apical channel contains primary human alveolar epithelial cells (ATI and ATII) with or without human CD14^+^ peripheral blood-derived macrophages, while the basal channel is lined by human primary pulmonary microvascular endothelial cells (HMVECs). **C.** Stitched image of the apical channel of the ALoC at 15 days of culture highlighting AT1 cells (HT1^+^, magenta) and AT2 cells (HT2^+^, green). Nuclei are labelled with DAPI (blue). Scale bar = 100 µm. **D (left).** Epifluorescent image of differentiated AT1 and AT2 cells on the ALoC. Scale bar = 50 µm. **D (right).** Three-dimensional immunofluorescence reconstruction of ALoC after 15 days of culture, highlighting differentiated AT2 cells and AT1 cells. Scale bar = 100 µm**. E.** Cross-sectional image of the ALoC highlighting AT1 and AT2 cells in a monolayer. Scale bar = 50 µm. Images are representative of at least 3 independent chips from 2 independent experiments.

Constructing a functional ALoC requires meticulously following a stepwise protocol (**Supplemental Figure 1A**). Briefly, the membranes of individual chips are chemically activated, coated with extracellular matrix and then human alveolar epithelial cells are added to the air (or apical) channel. After three days, human primary lung microvascular endothelial cells are introduced to the vascular (or basal) channel. Once cells are confluent (∼1 day), the alveolus-on-a-chip is connected to regulated media flow (30 uL/hr) in both channels. The next day, the apical channel media is removed, generating an air-liquid interface (ALI), and after 2 days of ALI, stretch is introduced to mimic breathing (5% stretch intensity, 0.2 Hz) and induce surfactant production (Thacker et al., 2020). After two days of stretch we add differentiated human CD14^+^ peripheral blood-derived macrophages (hereafter called ‘macrophages’) from anonymous donors to the top channel. To assess the donor compatibility of the alveolar epithelial cells and macrophages in the apical channel, we used immunofluorescence to compare AT1 and AT2 cell differentiation, cell morphology and distribution of cells along the apical channel of ALoCs with and without macrophages. In ALoCs without macrophages, a monolayer of both AT1 and AT2 cells was distributed equally across the apical channel (**Figure 1C, Supplemental Figure 1B**). High resolution images of ALoCs without macrophages highlighted distinct AT1 and AT2 cell types after 15 days in culture on the ALoC (**Figure 1D**). Three-dimensional reconstruction of the ALoC demonstrated the more cuboidal morphology of AT2 cells and the flatter morphology of AT1 cells (**Figure 1D, far right**). Cross sectional imaging of the airway channel further highlighted the nuclei in the basal region of the apical channel with interspersed AT1 and AT2 cells (**Figure 1E**). While the majority of cells were either AT1 or AT2 cells, a small number of KRT5 positive basal like cells were also observed in the airway channel (**Supplemental Figure 1C**). Finally, in the vascular channel, we observed a confluent monolayer of endothelial cells lining the entirety of the channel via staining for VE-Cadherin, a marker of endothelial cells (**Supplemental Figure 1D**).

We next turned to our analysis of ALoCs with macrophages (**Figure 2A**). In ALoCs with macrophages, the monolayer of differentiated AT1 and AT2 cells appears intact and unaltered by the addition of macrophages 24 hours prior (**Figure 2B, 2C, Supplemental Figure 1E**). Macrophages were observed residing above of the alveolar epithelial monolayer (**Figure 2D**). Although we introduced the same number of macrophages as alveolar epithelial cells, we observed many fewer macrophages overall, suggesting that the majority of added macrophages did not adhere tightly to the epithelial layer and were washed away (**Supplemental Figure 1F**). To address this question, we quantified the number of macrophages residing on three ALoCs and compared it to the number of macrophages seeded. We found that approximately 2.1x10^3^ total macrophages or 7% of the number seeded were present on the ALoCs at the end of the experiment. Stitched images of ALoCs containing macrophages also showed that, despite the loss of introduced macrophages, adherent macrophages were sufficient to evenly distribute across the entire apical channel (**Figure 2B; Supplemental Figure 1E**). Despite the addition of macrophages, AT cell distribution was comparable to uninfected ALoCs without macrophages (**Figure 1D**; **Figure 2C).** From these experimental observations, namely that AT1 and AT2 cell differentiation was maintained after the addition of primary human CD14^+^ peripheral blood macrophages, we concluded that macrophages could safely be added to the human ALoC model without dramatically altering epithelial biology.

**Figure 2.**
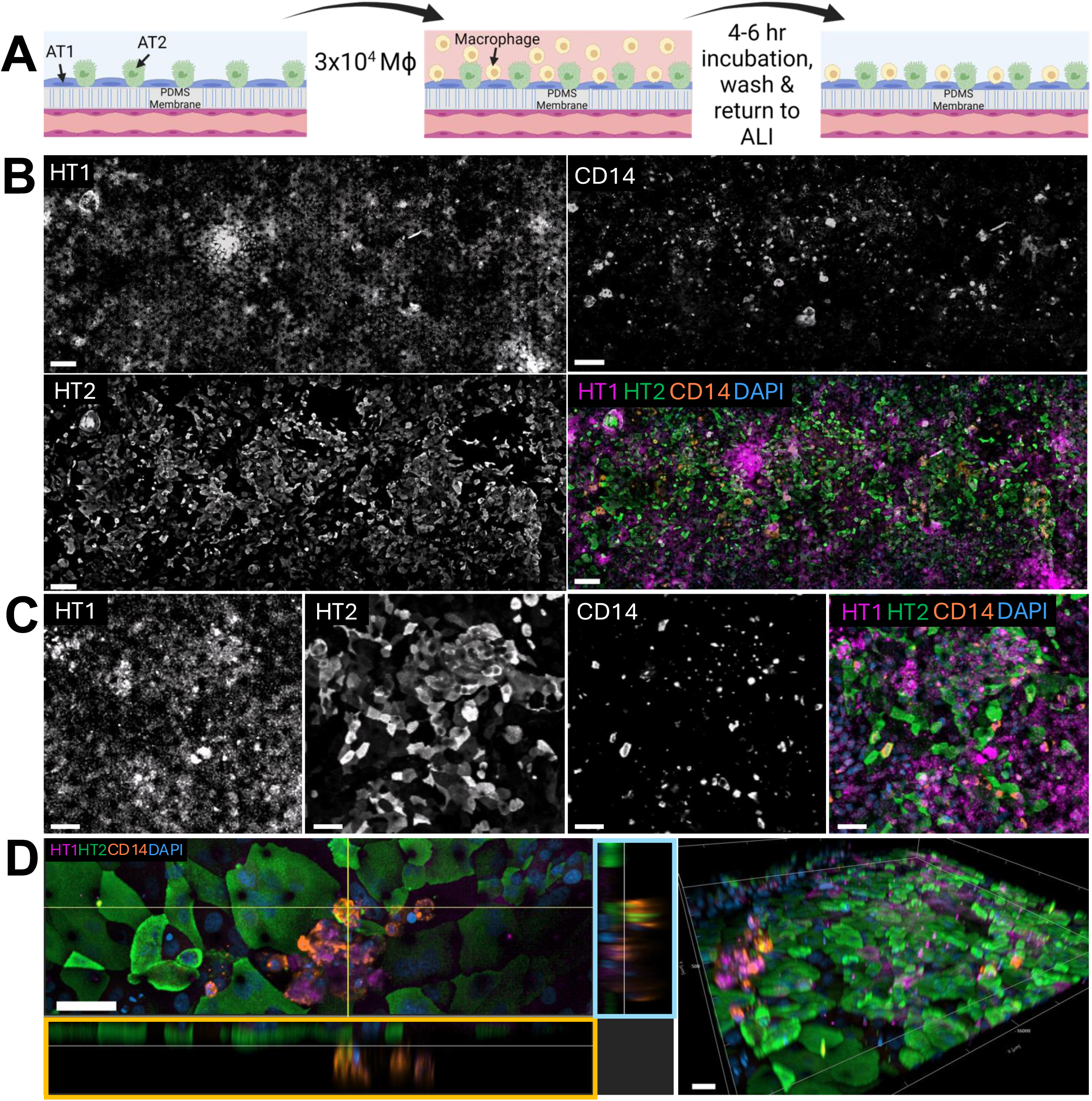
Human macrophages reside on the alveolar surface in the humanized ALoC. **A.** Schematic of the addition of macrophages to the apical channel. **B.** Stitched image of the apical epithelial channel of the ALoC after the addition of macrophages (CD14^+^, orange) on the ALoC. Nuclei are labelled with DAPI. Scale bar = 100 µm**. C.** Epifluorescent image of differentiated macrophages dispersed amongst AT1 and AT2 cells on the ALoC. Scale bar = 50 µm. **D (left).** Cross-sectional image of the ALoC showing AT1 cells (HT1^+^, magenta), AT2 cells (HT2^+^, green), and macrophages (CD14^+^, orange). Scale bar = 50 µm**. D (right).** Three-dimensional reconstruction of ALoC highlighting the addition of macrophages on the surface of differentiated AT1 and AT2 cells. Scale bar = 50 µm. Images are representative of at least 3 independent chips from 2 independent experiments. **E.** Cross-sectional image of the ALoC with macrophages highlighting macrophages laying on the surface of AT1 and AT2 cells. Scale bar = 50 µm. Images are representative of at least 3 independent chips from 2 independent experiments.

After establishing the ALoC model with human macrophages, we next incorporated mCherry-expressing *M. fortuitum* to the system. We introduced mCherry-expressing *M. fortuitum* at an estimated multiplicity of infection (MOI) of ∼1 for the approximate number of AT1 and AT2 cells on the chip (∼3.0 x 10^4^) (**Figure 3A**). We plated serial dilutions of the inoculum and determined the concentration of *M. fortuitum* added to the ALoCs was approximately 3.15x10^4^ CFU/mL. When ALoCs were infected with *M. fortuitum* in the absence of macrophages, we observed large clumps of bacteria scattered across the ALoC (**Figure 3B, 3C**). Along the apical channel in *M. fortuitum*-infected ALoC, we also observed areas that lacked strong immunofluorescence signal that we did not observe in uninfected ALoCs with or without macrophages (**Figure 3B, image center**). Higher resolution imaging analysis suggested loss of viability of AT1 and AT2 cells and a more rounded cell morphology, particularly for AT2 cells (**Figure 3C**). HT1 staining was also more diffuse in *M. fortuitum*-infected ALoCs. In addition, while ∼17% of cells were double positive for HT1 and HT2 staining in uninfected ALoCs (**Figure 3D, 3E**), we observed a significant increase in cells that stained with both HT1 and HT2 antibodies in *M. fortuitum*-infected ALoCs without macrophages (**Figure 3F, 3G)**. Though we observed large clumps of bacteria, *M. fortuitum* cord-like structures were not observed within the initial 24 hours of infection. Taken together, introducing *M. fortuitum* to the ALoC model at an approximate MOI of 1 generated a productive infection with sustained growth of bacteria and epithelial cell responses consistent with active infection.

**Figure 3.**
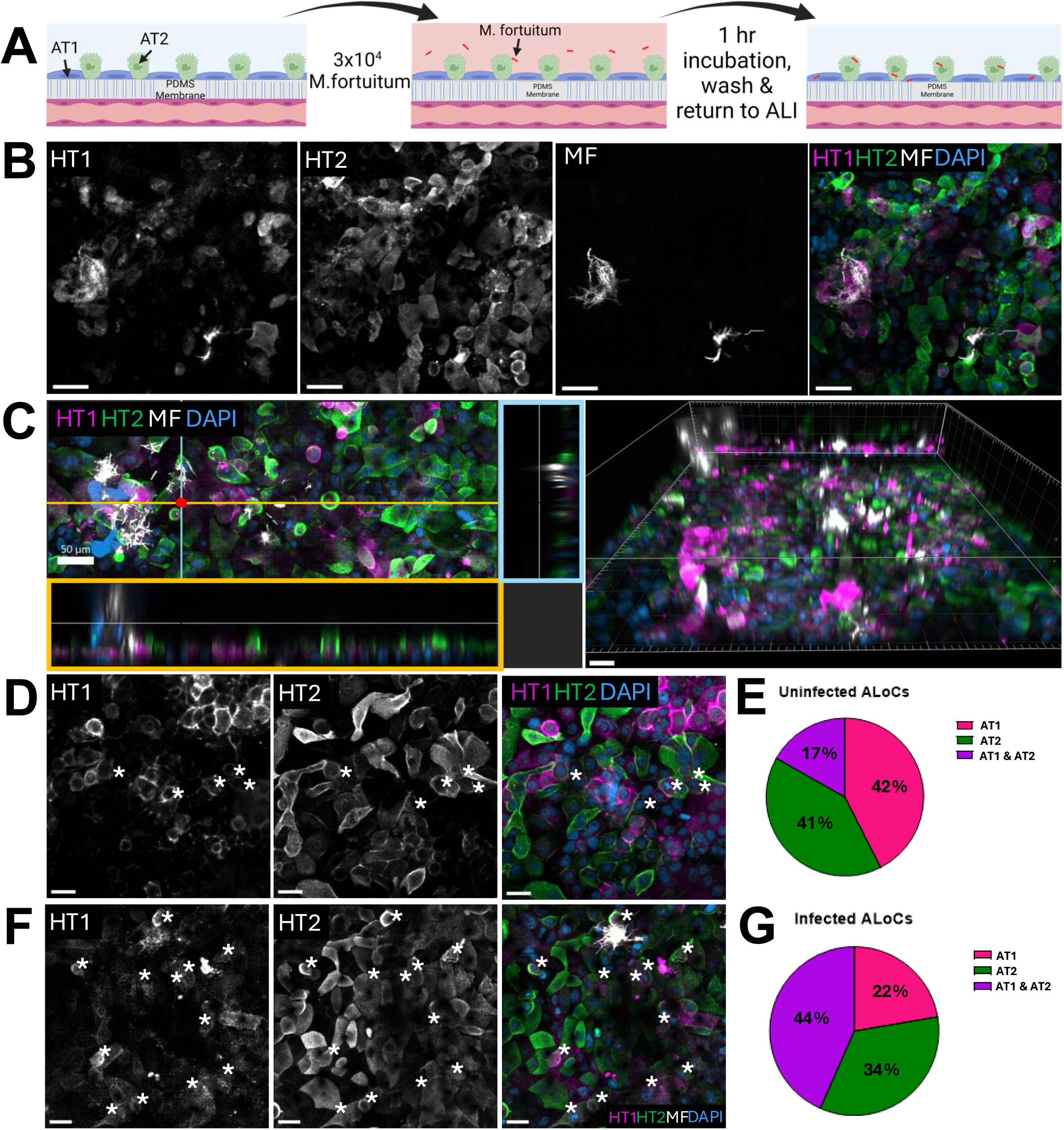
*Mycobacterium fortuitum* infection on an Alveolus Lung-on-a-Chip without macrophages. **A.** Schematic of *M. fortuitum* addition to the ALoC. mCherry-expressing *M. fortuitum* was added to the ALoC at an MOI of 1 with respect to the alveolar epithelial cells. After 1 hour of static incubation at 37°C, ALoCs were washed and returned to ALI. **B.** Epifluorescent image *M. fortuitum* (mCherry, white)-infected AT1 (HT1+, magenta) and AT2 (HT2+, green) cells. Scale bar = 50 µm. **C (left).** Cross-sectional image of *M. fortuitum* (mCherry, white) infection on ALoC with AT1 and AT2 cells. Scale bar = 50 µm**. C (right).** Three-dimensional reconstruction of an infected ALoC without macrophages. Scale bar = 50 µm**. D.** A single z-slice image of an uninfected ALoC without macrophages containing AT1, AT2 and AT1/AT2-positive cells, with AT1/AT2 cells highlighted by asterisks. Nuclei are labelled with DAPI (blue). Scale bar = 30 µm. E. Pie chart representing the ratio of AT1, AT2 or AT1/AT2-positive cells on ALoCs without macrophages. Cells were counted from 3 independent uninfected ALoCs without macrophages (number of cells counted per chip range from 900-1300 total cells) and averaged. **F.** A single z-slice image of an infected ALoC without macrophages containing AT1, AT2 and AT1/AT2-positive cells, with AT1/AT2 cells highlighted by asterisks. Nuclei are labelled with DAPI (blue).Scale bar = 30 µm. **G.** Pie chart representing the ratio of AT1, AT2 or AT1/AT2-positive cells on *M. fortuitum*-infected ALoCs without macrophages. Cells were counted from 3 independent infected ALoCs without macrophages (number of cells counted per chip range from 800-900 total cells) and averaged. Images are representative of at least 3 independent chips from 2 independent experiments.

We next wanted to determine how *M. fortuitum* grows on ALoC in the presence of human macrophages (**Figure 4A**). Twenty-four hours after seeding, ALoCs previously seeded with macrophages were infected with *M. fortuitum* as described above. Notably, the alveolar epithelial and pulmonary microvascular endothelial cells lining the chips were from the same donors as for ALoCs without macrophages. On ALoCs with macrophages infected with *M. fortuitum*, there was suitable distribution of AT2 cells across the chips (comparable to infected ALoCs without macrophages). In contrast to *M. fortuitum*-infected ALoC without macrophages, we observed that most bacteria were found to be intracellular within macrophages with significantly fewer bacterial clumps (**Figure 4B-D**) and cord-like structures were not observed. Cross-sectional and three-dimensional analysis of the ALoC with macrophages highlighted that *M. fortuitum* were residing within macrophages (**Figure 4D**). To quantify bacteria within either AT1 or CD14+ cells, we analyzed 10 z-stacks from 3 independent ALoCs containing macrophages. We used three-dimensional reconstructions and ortho-projections to assign bacteria within AT1 cells (HT1+) or macrophages (CD14^+^) and found that bacteria were found more often in macrophages than AT1 cells (**Figures 4B, 4E)** To quantify bacteria within either AT2 cells (HT2+) or macrophages (CD14^+^), we performed a similar analysis with 3 ALoCs containing macrophages and determined that bacteria were found more often in macrophages than AT2 cells (**Figures 4C, 4F**). Of note, due to technical limitations of fluorophores we could use, quantifying bacteria within AT1 cells, AT2 cells, and macrophages simultaneously could not be done within the same AloC. Taken together, we conclude that in the presence of macrophages overlying alveolar epithelial cells, *M. fortuitum* are ingested by macrophages, preventing extracellular growth and bacterial clumping. Such an infection may also prevent loss of epithelial cell integrity.

**Figure 4.**
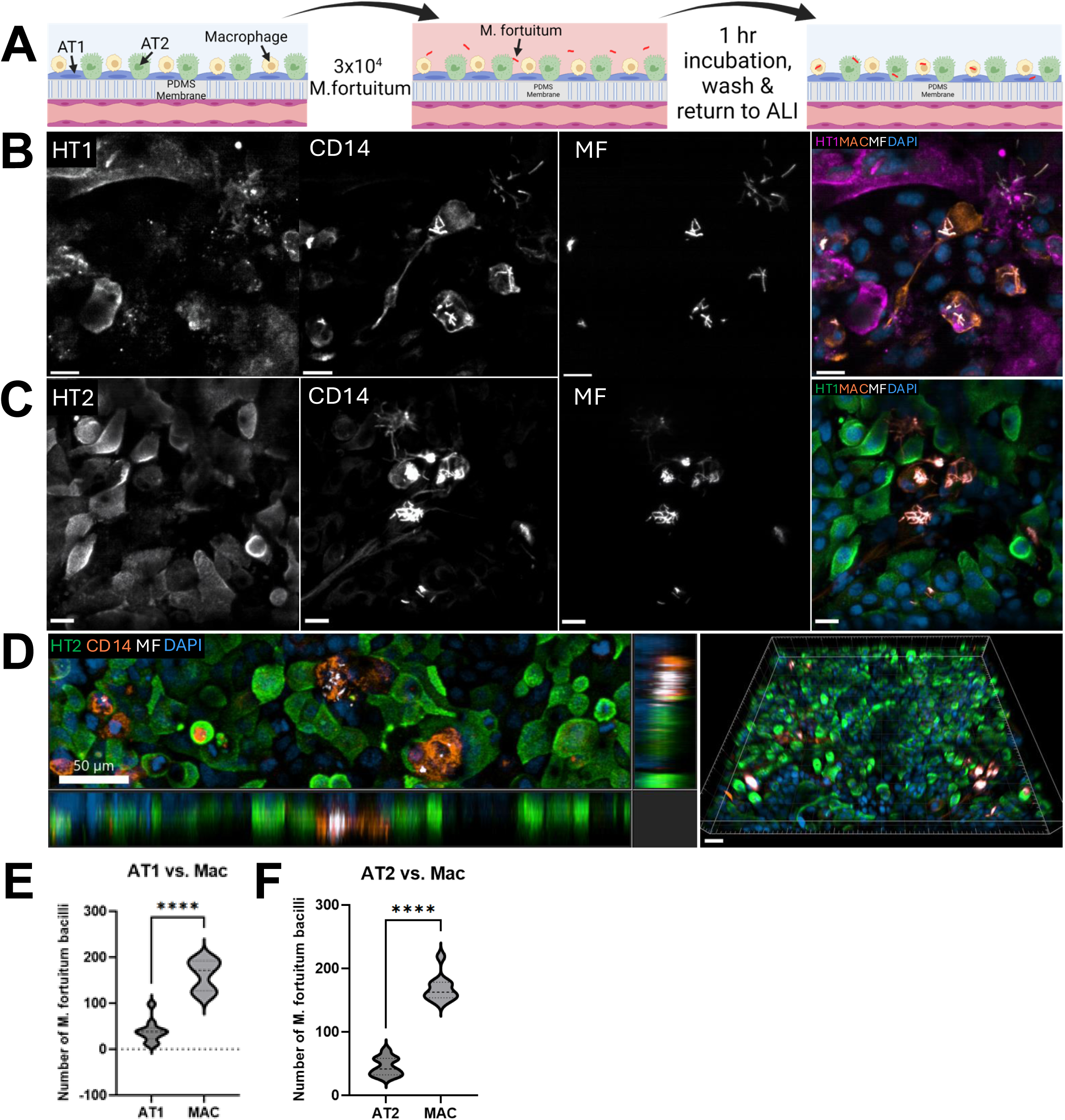
*Mycobacterium fortuitum* infection on an Alveolus Lung-on-a-Chip containing macrophages. **A.** Schematic of *M. fortuitum* infection on ALoC containing macrophages. 24 hours after the addition macrophages, *M. fortuitum* was added to the ALoC at an MOI of 1 with respect to the alveolar epithelial cells, incubated at 37°C for 1 hour, washed, and returned to ALI. **B**. Epifluorescent image of an *M.fortuitum* (mCherry, white)-infected ALoC with macrophages containing AT1 cells (HT1+, magenta) and macrophages (CD14+, orange). Scale bar = 20 µm. **C.** Epifluorescent image of an *M.fortuitum* (mCherry, white)-infected ALoC with macrophages containing AT2 cells (HT2+, green) and macrophages (CD14+, orange). Scale bar = 20 µm. **D (left).** Cross-sectional image of *M. fortuitum* infection on ALoC with the addition of macrophages, highlighting intracellular M. fortuitum bacilli within macrophages (CD14+, orange) rather than AT2 (HT2+, green) cells. Scale bar = 50 µm. **D (right).** Three-dimensional reconstructed image of an ALoC with macrophages containing AT2 cells and *M.fortuitum*-infected macrophages. Scale bar = 100 µm. **E.** Quantification of *M. fortuitum* bacilli found intracellularly within macrophages or AT1 cells. Each point represents the cumulative number of *M. fortuitum* bacilli counted in each cell type for 10 fields within a single chip (N=3 independent chips, p<0.0001 by unpaired Student’s t-test). **F.** Quantification of *M. fortuitum* bacilli found intracellularly within macrophages or AT2 cells. Each point represents the cumulative number of *M. fortuitum* bacilli counted in each cell type for 10 fields within a single chip (N=3 independent chips, p<0.0001 by unpaired Student’s t-test). Images are representative of at least 3 independent chips from three independent experiments.

To assess the response of human ALoC to *M. fortuitum* infection, we used bulk RNA sequencing to compare the transcriptional status of *M. fortuitum* infected ALoC with macrophages to uninfected ALoC with macrophages. After infection at an MOI of 1 for 24 hours, we collected total RNA from the apical channel containing AT1 cells, AT2 cells and macrophages for sequencing. While there was some heterogeneity in the response, as would be expected in a primary infection, we observed 1454 genes were more than 2-fold upregulated, while 588 were more than 2-fold downregulated (**Figure 5A, B**). Amongst the upregulated genes, we observed significant upregulation of key cytokines such as tumor necrosis factor (*TNF*), granulocyte-macrophage colony-stimulating factor (*GM-CSF* or *CSF2*), macrophage colony-stimulating factor (*M-CSF* or *CSF3*), interleukin 1A (*IL1A*), interleukin 1B (*IL1B*), interleukin 6 (*IL6*), and interleukin 8 (*IL8*), along with the alarmin, calprotectin (*S100A8* and *S100A9*) (**Figure 5C**). We also observed several chemokines upregulated including the chemokine (C-X-C motif) ligand family members *CXCL1, CXCL2, CXCL3, CXCL5, CXCL6, CXCL10* and *CXCL11* and the C-C motif chemokine ligand family members *CCL2, CCL5, CCL20* and *CCL28* (**Figure 5C**). Finally, we also noted that many secreted serine protease inhibitor (SERPIN) genes were also upregulated by *M. fortuitum* infection (**Figure 5D**). Using pathway analysis (Ge et al., 2018), we found that the most significantly upregulated processes were involved in host defense including “response to bacterium”, “defense response to bacterium”, “antimicrobial humoral response”, “response to molecule of bacterial origin” and “response to lipopolysaccharide” (**Table 1**). Likewise, upregulated molecular functions included several annotated as involved in “signaling” in addition to “cytokine” and various channel functions (**Table 2**). Taken together, our data highlight the acute inflammatory response of humanized ALoC containing human macrophages to *M. fortuitum* infection.

**Figure 5.**
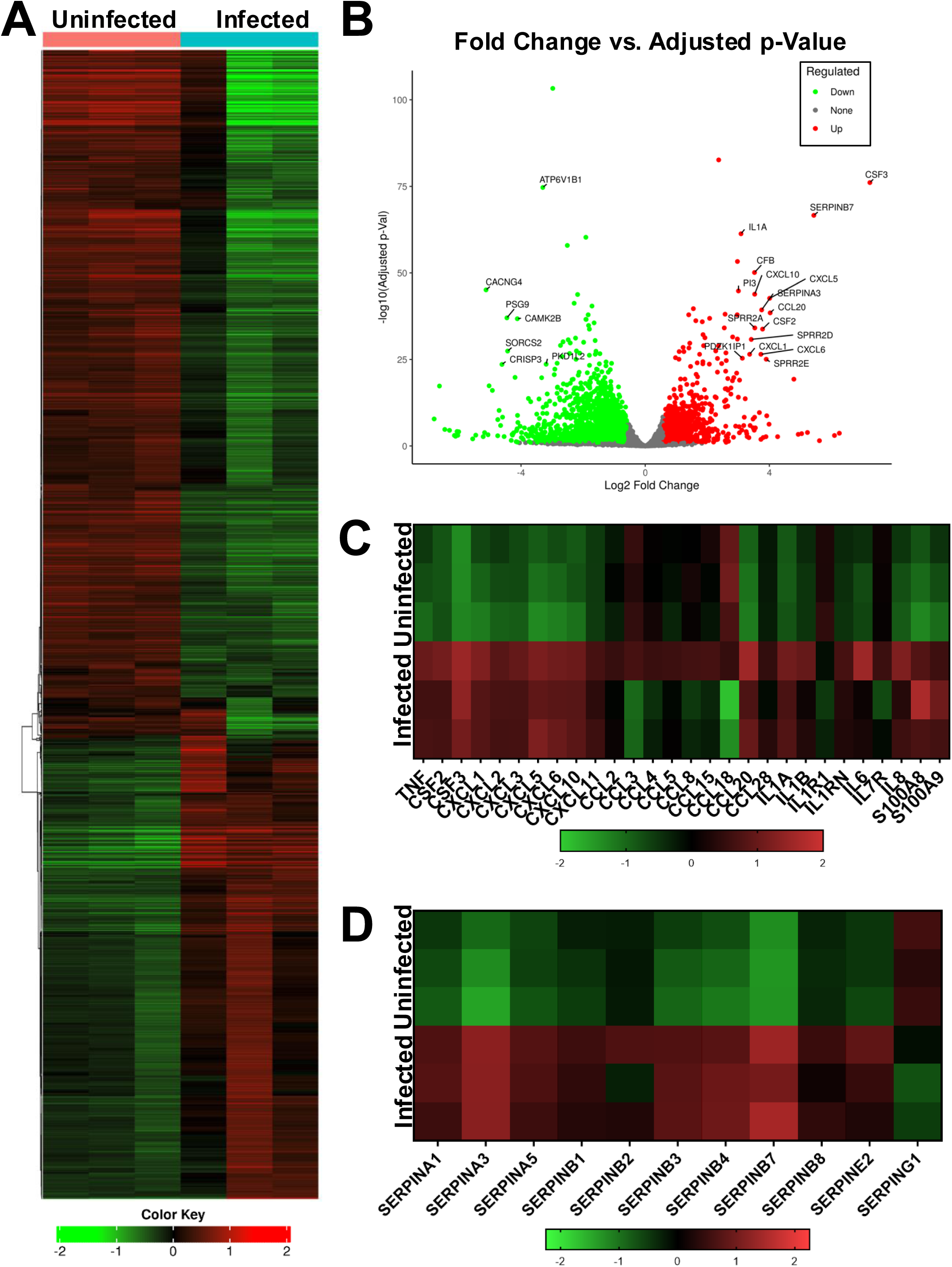
Bulk RNA-seq Analysis of *M. fortuitum*-infected ALoCs with macrophages. **A.** Heat-map of the top 2000 genes expressed in uninfected and *M. fortuitum*-infected ALoCs (24 hours post-infection) (N=3 per group) using Pearson distance and average linkage. **B.** The -log_10_ (Adjusted p-Value) was plotted against log_2_ fold change to create a volcano plot representing the top 25 differentially expressed genes (DEGs). Genes with an adjusted p-value<25 and log_2_ fold change<2 were filtered from this analysis. **C.** Heat map comparing the log_2_ fold change of the most differentially expressed chemokines and cytokines in ALoCs infected with *M. fortuitum* versus uninfected ALoCs after 24 hours of infection. **D.** Heat map comparing the log_2_ fold change of the most differentially expressed SERPINs in ALoCS infected with *M. fortuitum* versus uninfected ALoCs after 24 hours of infection.

**Table 1.** The most significant up-regulated and down-regulated GO Biological Process pathways from M. fortuitum-infected ALoCs containing macrophages.

| Direction | GO Biological Process | Statistic | Genes | adj.Pval |
| --- | --- | --- | --- | --- |
| Up | <u>Response to bacterium</u> | 7.3996 | 438 | 1.1e-09 |
|  | <u>Humoral immune response</u> | 6.5518 | 138 | 6.0e-07 |
|  | <u>Defense response to bacterium</u> | 6.3097 | 169 | 1.1e-06 |
|  | <u>Antimicrobial humoral response</u> | 6.0602 | 65 | 1.9e-05 |
|  | <u>Response to molecule of bacterial origin</u> | 5.5061 | 251 | 3.7e-05 |
|  | <u>Chemotaxis</u> | 5.2453 | 444 | 8.2e-05 |
|  | <u>Taxis</u> | 5.2402 | 445 | 8.2e-05 |
|  | <u>Response to lipopolysaccharide</u> | 5.2654 | 240 | 8.2e-05 |
|  | <u>Antimicrobial humoral immune response mediated by antimicrobial peptide</u> | 5.6504 | 41 | 1.4e-04 |
| Down | <u>MRNA processing</u> | -6.8939 | 402 | 4.4e-08 |
|  | <u>RNA splicing</u> | -6.5273 | 385 | 2.3e-07 |
|  | <u>Histone modification</u> | -5.7112 | 429 | 1.6e-05 |
|  | <u>Proteasome-mediated ubiquitin-dependent protein catabolic process</u> | -5.5755 | 410 | 2.5e-05 |
|  | <u>Proteasomal protein catabolic process</u> | -5.534 | 470 | 2.5e-05 |
|  | <u>Chromatin organization</u> | -5.4523 | 478 | 3.2e-05 |
|  | <u>NcRNA metabolic process</u> | -5.3774 | 477 | 4.2e-05 |
|  | <u>Peptidyl-lysine modification</u> | -5.128 | 343 | 1.5e-04 |
|  | <u>RNA splicing via transesterification reactions with bulged adenosine as nucleophile</u> | -5.1146 | 265 | 1.5e-04 |
|  | <u>MRNA splicing via spliceosome</u> | -5.1146 | 265 | 1.5e-04 |
|  | <u>RNA splicing via transesterification reactions</u> | -5.097 | 269 | 1.5e-04 |

**Table 2.**
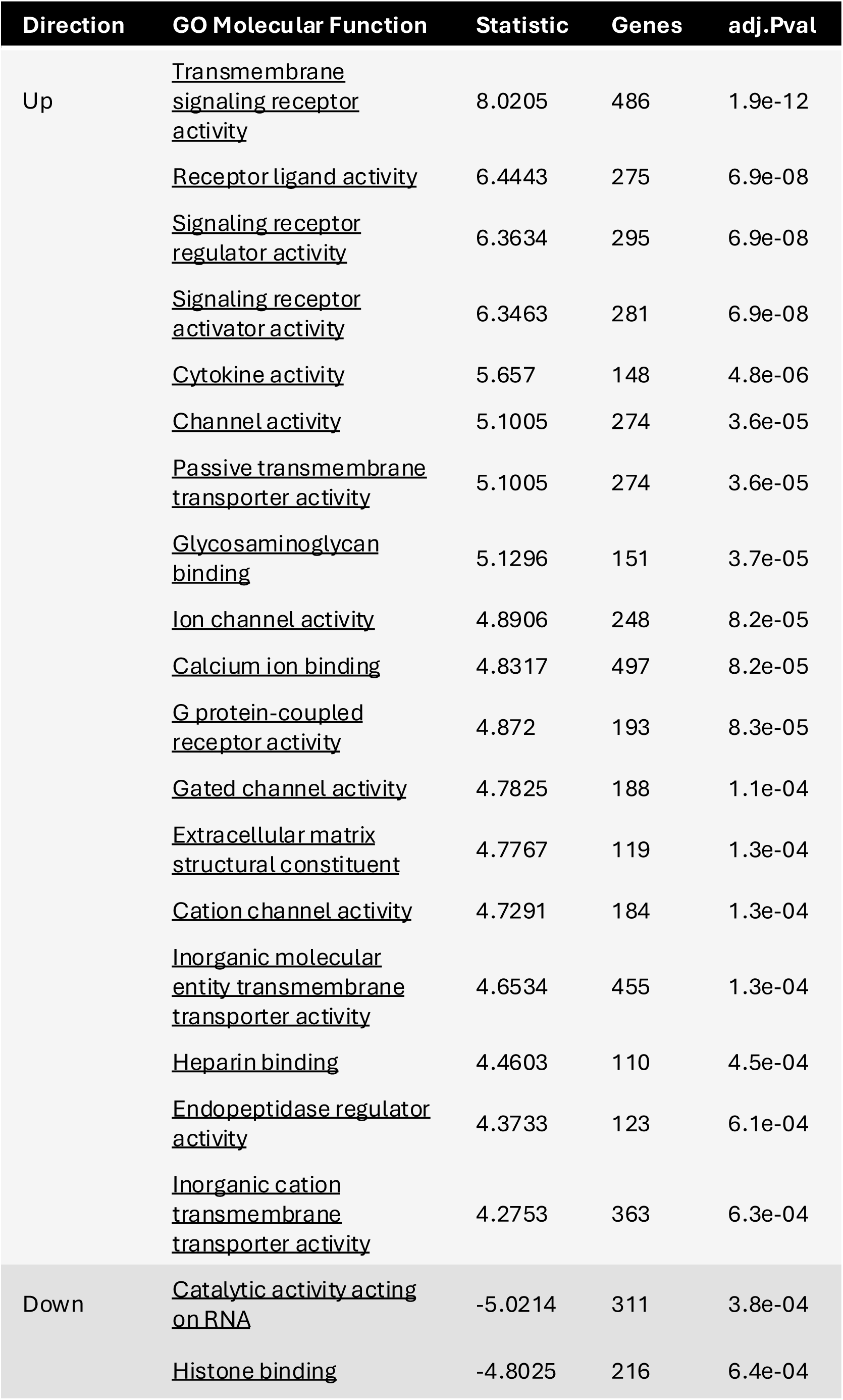
The most significant up-regulated and down-regulated GO Molecular Function pathways from M. fortuitum-infected ALoCs containing macrophages.

## Discussion

We report the development of a fully humanized ALoC model for alveolar NTM infection. This model develops features consistent with initial primary alveolar infection, including infection of both airway macrophages and occasional alveolar cells. Moreover, bulk RNA sequencing reveals activation of an acute inflammatory signaling cascade, including induction of cytokines, chemokines and alarmins like serpins and calprotectin, with potential roles in recruitment of myeloid and lymphoid cells from the vasculature and in triggering a robust immune response to restrict bacterial growth. These findings highlight the value of using the humanized lung on a chip to model a primarily human infectious disease.

Prior studies using the alveolus-on-a-chip have included infection models, such as infection with *E. coli* (Huh et al., 2010), *M. tuberculosis* (Thacker et al., 2020), *Aspergillus fumigatus* (Hoang et al., 2022), *Staphylococcus aureus* (Bai et al., 2022, Deinhardt-Emmer et al., 2020), and influenza virus (Bai et al., 2022, Deinhardt-Emmer et al., 2020). The alveolus-on-a-chip has also been used to determine the effect of mechanical strain on inflammation and particle uptake (Huh et al., 2010), pulmonary thrombosis (Jain et al., 2018), and mechanotransduction and pulmonary edema (Li et al., 2019). In the context of *M. tuberculosis* and *A. fumigatus* infection models, addition of human monocytes/macrophages impacts pathogenesis (Thacker et al., 2020, Hoang et al., 2022). With *M. tuberculosis*, cord-like structures are observed compressing macrophage nuclei (Mishra et al., 2023). We did not observe a similar phenomenon with *M. fortuitum* infection of human macrophages on the ALoC, which might be due to differences in the cell walls of the organisms or other factors that impact the growth characteristics of *M. tuberculosis* (slow) as compared to *M. fortuitum* (rapid). Alternatively, differences in the time after infection used for analysis (24 hours in this study versus 72 hours (Mishra et al., 2023)) or imaging techniques may also account for these differences. In a different model system using human bronchial epithelial cells, some mycobacteria formed surface biofilms that ultimately caused apical non-apoptotic cell death (Barclay et al., 2024). Because our infection model was limited to 24 hours, we did not observe biofilm formation, but we did note some loss of cell viability and monolayer disruption, consistent with the observations made with other mycobacterial species (Barclay et al., 2024).

We identified a transcriptional signature of early human alveolar infection with *M. fortuitum*. Some notable findings include marked upregulation of important cytokines and chemokines that may be involved in stimulating an early innate immune response to airway infection. For example, both IL6 and IL8 are key cytokines involved in neutrophil recruitment to the airway (Fu and Harrison, 2021), and such recruited cells could have both beneficial (contributing to NTM killing) and detrimental (involvement in tissue damage and bronchiectasis) effects (Alkarni et al., 2023). While there have been previous studies of human peripheral blood transcriptional responses in the context of pulmonary NTM infection (Wang et al., 2024, Cowman et al., 2018, Lindestam Arlehamn et al., 2022) or tuberculosis (Tabone et al., 2021, Berry et al., 2010), as well as efforts to determine transcriptional responses in human lung tissue from tuberculosis patients (Wang et al., 2023a), to our knowledge this is the first analysis of the early interaction of an NTM with a humanized ALoC. Interestingly, in the RNA sequencing analysis of PBMCs from individuals infected with *M. avium*, TNF signaling was enriched, and individual genes such as those for *SERPINA1* and calprotectin were modestly upregulated (Lindestam Arlehamn et al., 2022). Calprotectin was also upregulated by *M. avium* infection in an air-liquid interface model of primary human bronchial epithelial cells (Barclay et al., 2023). Likewise, in a proteomics analysis of serum exosomes in humans with NTM infection, a variety of serpins including SERPINA1 and SERPINA5 were enriched in patients with *M. abscessus* or *M. avium* infection (Wang et al., 2023b). Recent work has found the induction of cytosolic SERPINs may protect *M. marinum* infected macrophages from cathepsin B mediated lysosomal rupture and cell death (Nobs et al., 2024), a process called lysoptosis that is evolutionary conserved from *C. elegans* to mammals (Luke et al., 2022). Thus, SERPIN induction by *M. fortuitum* infection, potentially within infected macrophages, may act to protect infected cells from lysoptosis. Of note, because we used bulk RNA sequencing, we were unable to identify the precise cellular source of transcriptional changes identified in our analysis. Future single cell sequencing studies will be necessary to determine the responses of individual cell types within the model system, and how such responses impact intercellular communication. Taken together, our bulk RNAseq analysis has identified unique features of early alveolar NTM infection.

There are several limitations to our model. First, because our model uses all primary human cells, genetic manipulation of the various cellular constituents is impractical, though primary human monocytes can be transduced with lentivirus to abrogate gene expression (Campbell and Spector, 2012, Franco et al., 2017). Alternatively, use of genetically modified human or mouse cell lines can overcome this limitation, though immortalized cell lines have their own inherent drawbacks. Second, because primary cells, whether pulmonary epithelial cells, pulmonary vascular cells or PBMCs, are obtained from unique individuals, genetic variation generates greater complexity. Third, to optimize infection and cognizant that not all bacteria would adhere to the alveolar channel or be ingested by macrophages or AT1/AT2 cells, we chose to use an MOI of 1, which is not likely to be reflective of the number of bacteria that reach the terminal alveoli during a natural infection. Future work will compare a range of MOI with the ALoC system and other NTM. Finally, though our model does allow multiple cell types (epithelia, endothelia and myeloid cells) to interact with each other and the NTM in the alveolar microenvironment, the interactions are limited to the cells added to the chamber, and thus do not include all the possible myeloid and lymphoid cells that could be recruited to the airway in the setting of pulmonary mycobacterial disease, such as neutrophils (Kimmey et al., 2015), eosinophils (Bohrer et al., 2021) or innate lymphoid cells (Ardain et al., 2019).

In conclusion, we have developed a fully humanized alveolus on a chip model of pulmonary NTM infection. By capturing the complexity of NTM in three dimensions and with multiple cell types, as would be found in a human lung, this model creates an opportunity to better characterize the cellular and molecular mechanisms that mediate the outcome of human airway NTM infections. Because NTM infections are on the rise, particularly in individuals with cystic fibrosis, bronchiectasis and people with HIV, it may be useful to apply this technology to determine the relative impact of genetic risk alleles (i.e. for those with cystic fibrosis) or coinfection with HIV on the alveolar transcriptional responses or bacterial survival in the alveolus over time. Furthermore, by leveraging this system to directly compare responses to other common human pulmonary NTM infections, such as *M. avium* (Cristancho-Rojas et al., 2023, Varley and Winthrop, 2022, Donohue, 2021), *M. kansasii* and *M. abscessus* (Toure et al., 2023, Lagune et al., 2023, Cristancho-Rojas et al., 2023, Ordway et al., 2008, Klever et al., 2023) future studies could identify shared and unique survival strategies used by each organism, and establish distinct immune pathways responsive to each pathogen which could function to limit or exacerbate infection, or be used clinically as biomarkers of disease. Finally, as new drugs are developed to treat pulmonary NTM infection, we envision applying this model for the simultaneous assessment of efficacy (i.e. reduction in bacterial load) and cellular toxicity via infusion of test compounds into the vascular channel.

## Materials and Methods

### Bacteria

*Mycobacterium fortuitum* subsp. *fortuitum* strain was obtained from ATCC (strain TMC 1529). We transformed *M. fortuitum* with a vector containing *mCherry* driven by the GroEL constitutive promoter.

### Primary Cell Culture

Primary human alveolar epithelial cells (ATs) were obtained from Cell Biologics (H-6053). Prior to seeding on Alveolus Lung-on-Chips (ALoCs), ATs were expanded *in vitro* in a T25 flask coated with a 1% gelatin-based coating solution (Cell Biologics, catalog #6950) and complete medium comprising of base media and supplements (Lonza, CC-3118). Medium was prepared according to manufacturer’s instructions using all supplements except GA-1000 where 1% Pen-Strep solution (Gibco 15140-122) was added instead along with 5% FBS and hereby called Small Airway Growth Medium (SAGM). Primary human microvascular endothelial cells were purchased from Lonza (CC-2527). Human Lung Microvascular Endothelial cells (HMVECs) were expanded in a T75 in complete medium comprising of base media and supplements (Lonza, CC-3202). Medium was prepared according to manufacturer’s instructions using all supplements except GA-1000 where 1% Pen-Strep solution was used instead along with 5% FBS and hereby called EGM-2MV. Both primary cell lines were cultured at 37°C in 5% CO_2_ until ∼80% confluency before detachment with TrypLE Express (Gibco 12604013) and use on the ALoC.

One week prior to seeding the macrophages on the ALoC, peripheral blood mononuclear cells (PBMCs) were obtained from buffy coats from anonymous donors (Carter BloodCare). PBMCs were isolated using Ficoll (Cytivia, 17144003) and SepMate50 tubes from StemCell Technologies (85450). CD14^+^ monocytes were positively selected from the PBMCs using CD14 Microbeads (Miltenyi, 130-050-201) and seeded onto uncoated petri dishes. The monocytes were cultured overnight in RPMI medium (Gibco 11875-093) supplemented with 10% heat-inactivated human serum obtained from the buffy coat of the donor, 1% HEPES buffered solution (Lonza CC-5022), 1% sodium pyruvate (Gibco 11360-070), and 50 ng/mL human granulocyte-macrophage colony-stimulating factor (GM-CSF) (Peprotech 300-03-100UG) and is hereby called macrophage medium. The next day, the serum in the medium was changed to 10% FBS for the remainder of the culturing. CD14^+^ monocytes were differentiated to human monocyte-derived macrophages (HMDMs) for 7 days at 37°C in 5% CO_2_ until use on the ALoCs. GM-CSF was added to the macrophage medium for the first 4 days to differentiate the CD14^+^ monocytes.

### Human Alveolus LoC Model

ALoCs fabricated with polydimethylsiloxane (PDMS) were purchased from Emulate. The chips were activated using a 0.5 mg/mL solution comprised of ER-1 and ER-2 (Emulate) that was protected from light during the activation process. Working in a dark biosafety cabinet, both the top and bottom channel were filled completely with ER-1/ER-2 and placed under a UV light for 10 min, inspected for bubbles with a brightfield microscope and placed under the UV light for an additional 10 min. After activation of all chips, the top and bottom channels were coated with an extracellular matrix (ECM) solution specific to the cell type. The top channel (ATs) was coated with an ECM solution containing Collagen IV at 200 µg/mL (Sigma C5533), Fibronectin at 30 µg/mL (Millipore Sigma F2006) and Laminin at 5µg/mL (Sigma L6274). The bottom channel (endothelial cells) was coated with an ECM solution containing Collagen IV at 200 µg/mL and Fibronectin at 30 µg/mL. All ECM components and solutions were kept on ice and prepared by manufacturer’s recommendations. With an empty 200 µL pipette tip plugged into each outlet port, 100 uL of ECM solution were added to the respective channels by expelling ECM solution from pipette tip and plugging the inlet ports with the tip. The ECM-coated ALoCs were incubated at 4°C overnight and the next day each channel was washed twice with 200 µL of respective medium. Respective mediums were left in each channel prior to cell seeding and ALoCs were stored at 4°C until cell seeding.

ATs were seeded first in the top channel of all ALoCs. After detaching the cells from the T75 flasks, they were adjusted to a concentration of 1 x 10^6^ cells/mL in SAGM medium described above. The bottom channels were filled with SAGM medium during epithelial cell seeding. 50 µL of the AT cell suspension was pipetted into the top channel rapidly and checked under a microscope to ensure correct seeding density and cell homogeneity. Of note, the top channel volume is ∼28 µL total, but to ensure no bubbles are introduced, the channels are overfilled and the flow-through gently aspirated. Once all chips were seeded, they were placed in a chip cradle (Emulate) and incubated at 37°C for at least 2 hours. After confirming all cells had attached, each top and bottom channel were gently washed with 200 µL of warm SAGM medium and incubated overnight at 37°C. HMVECs were seeded 2-3 days later in the bottom channels of all ALoCs. Prior to HMVEC cell seeding, media was replenished daily for the ATs seeded on the ALoCs. The SAGM medium used for maintenance of the ATs on the chips (hereby called AT Maintenance Medium) was supplemented with Dexamethasone at 100 nM (Sigma D4902), Keratinocyte growth factor (KGF) at 5 ng/mL (Thermo Fisher PHG0094), 8-Br-cAMP at 50 µM (Sigma B7880), and Isobutyl methylxanthine (IBMX) at 25 µM (Sigma I7018). When HMVECs reached ∼80% confluency, they were detached and adjusted to a concentration of 5 x 10^6^ cells/mL in EGM-2MV medium (described above). The bottom channels of all ALoCs were filled with EGM-2MV medium prior to seeding. 20 µL of the HMVEC cell suspension was pipetted rapidly into the bottom channel and checked under the microscope for correct seeding density and cell homogeneity. The bottom channel volume is ∼6 µL total. Similarly to the top channel, the bottom channel was overfilled to prevent the introduction of bubbles to the channel. The chip was then immediately flipped upside down and placed in the chip cradle to ensure cells were seeded on the porous membrane in the bottom channel. Once all ALoCs were seeded and flipped upside down, they were incubated at 37°C for at least 2 hours until all cells were attached. After all cells had attached, the bottom channel was washed with 200 µL of warm EGM-2MV medium and incubated overnight at 37°C.

Prior to attaching ALoCs to Pods, warm EGM-2MV and AT Maintenance medium were degassed using a Steriflip device for 5 min each to prevent air bubbles being trapped in the microfluidic lines of the Pod. 3 mL of each medium were pipetted into its respective reservoir within the Pod. Using Emulate’s Zoe, the Pods were primed with medium to prevent bubbles from impeding medium flow to the ALoCs. Once primed, each ALoC was snapped into each respective Pod. To further help prevent bubble formation within the microfluidic lines, a Regulate Cycle was performed on each chip via the Zoe. The ALoCs from this point on were harbored within the Pods and fresh medium was exchanged through the chip at a flow rate of 30 µL/hr supplied by the Zoe.

After 24 hours of continuous media flow on all ALoCs, air-liquid interface (ALI) was introduced to all ALoCs by removing all medium from the top channel and changing the top channel from liquid media flow to air. The bottom channel medium was changed to ALI medium which comprised of a Medium 199 base (Thermo Fisher 11043023) supplemented with 10 ng/mL Human Epidermal Growth Factor (Peprotech AF-100-15), 3 ng/mL Human Basic Fibroblast Growth Factor (Peprotech AF-100-18B), 0.125 ng/mL Human Vascular Endothelial Growth Factor (Peprotech AF-100-20), 1 µg/mL Hydrocortisone (Sigma H0135), 10 µg/mL Heparin (Sigma H3149), 80 µM di-butyryl cAMP (Sigma B7880), 1mM L-Glutamax (Gibco 35050-061), 20 nM Dexamethasone (Sigma D4902), 1% Pen-Srep solution (Gibco 125140-122), and 2% FBS. Bottom channel liquid flow rate was set to 30 µL/hr and fresh ALI medium was replenished in the bottom channel reservoir every 2-3 days. After 48 hours of ALI, mechanical stretch (5%, 0.20 Hz) was initiated using the Zoe. All ALoCs were maintained in this way until seeding of macrophages and infection with *M. fortuitum*.

### Macrophage Seeding on ALoCs

After 7 days of differentiation with M-CSF to HMDMs, macrophages were detached from petri dishes using ice-cold 5 mM EDTA in PBS and gentle scraping. HMDMs were centrifuged at 300xg for 5 min and resuspended in a 1:1 ratio of AT Maintenance Medium (without dexamethasone) and Macrophage Medium to a concentration of 1 x 10^6^ cells/mL. ALoCs were removed from the Zoe and detached from the Pods, and top channel was washed with 200 µL of HPAEC Maintenance Medium. With medium still in the top channel, 50 µL of HMDM cell suspension was rapidly pipetted through the top channel. ALoCs were placed in square petri dishes and incubated at 37°C with 5% CO_2_ for 3-4 hours to allow attachment to the top channel. Once attached, ALoCs were attached back to Pods after priming and after 2 hours of a Regulate Cycle were returned to ALI.

### *M. fortuitum* Infection on Human ALoC

Upon returning ALoCs to ALI conditions after macrophage attachment to the ALoCs, ALI medium was replaced with ALI medium without penicillin/streptomycin, to avoid impacting subsequent bacterial infection. Twenty-four hours after returning to ALI conditions, the ALoCs were infected with *M. fortuitum*. mCherry-expressing *M. fortuitum* was cultured at 37°C in liquid 7H9 medium (BD Difco 271310) supplemented with 10% OADC enrichment (BD 212351), 50 µg/mL Kanamycin (Sigma 60615) and 0.05% Tyloxapol (Sigma T8761) until it reached an OD_600_ 0.5 – 0.7. The culture was pelleted by centrifugation at 3500 RPM for 10 min. The pellet was washed three times with 50 mL of 1X PBS (-Ca^+^/-Mg^+^) by centrifugation at 3500 RPM for 10 min, and resuspended in 5 mL PBS for a slow spin to remove cell debris at 500 RPM for 5 min. The supernatant was collected and passed through a 26-gauge needle three times to generate a single-cell suspension. The bacterial suspension was adjusted to an MOI of 1 with respect to the epithelial cells in HPAEC Maintenance Medium, and 50 µL of the *M. fortuitum* suspension was pipetted rapidly through the top channel. The ALoCs were incubated at 37°C for 1 hr. under static conditions, and then washed three times with epithelial medium before returning ALoCs back to Pods and into the Zoe for another Regulate Cycle. ALI was initiated again and the ALoCs were infected for a total of 24 hours.

### RNA Extraction from ALoCs

ALoCs were removed from the Zoe and Pods. Top and bottom channels were washed three times with 200 µL of ice-cold 1X DPBS (-Ca^+^/-Mg^+^). An empty p200 filtered tip was inserted into the bottom inlet, bottom outlet, and top outlet ports. 200 µL of TRIzol reagent (Invitrogen #15596026) was pipetted in each ALoC by rapidly pressing and releasing the plunger three times before collecting the supernatant in an RNase-free 1.5 mL tube. This process was repeated once more for a total of ∼400 µL of supernatant. TRIzol was added to each tube of RNA to a final volume of 1 mL. The RNA was stored at -80°C before extraction using the Qiagen RNeasy Mini Columns (74106). AT and macrophage lysates were thawed on ice and incubated at room temperature for 5 min. 200 µL of chloroform per 1 mL of lysates was added to each tube, shaken vigorously for 15 sec., incubated at room temperature for 2-3 min, and centrifuged for 5 min at 12,000xg at 4°C. The aqueous phase was transferred to fresh RNase-free tubes and the rest of the RNA extraction was performed according to the Qiagen RNeasy Mini Column Kit. Extracted RNA was stored at -80°C until use for library preparation and RNA-sequencing.

### Library Preparation and RNA-sequencing

Samples were analyzed on an Agilent Tapestation 4200 to determine level of degradation to ensure that only high quality RNA was used (RIN Score 8 or higher). We used a Qubit 4.0 Fluorimeter (ThermoFisher) to determine RNA concentration prior to starting library prep. One microgram of total DNAse treated RNA was then prepared with the TruSeq Stranded mRNA Library Prep Kit (Illumina). Poly-A RNA was purified and fragmented before strand-specific cDNA synthesis. cDNA was then A-tailed and indexed adapters ligated. After adapter ligation, samples were PCR amplified and purified with AmpureXP beads, then validated again on the Agilent Tapestation 4200. Before being normalized and pooled, samples were quantified by Qubit then sequenced on the Illumina NextSeq 2000 using a P2-100 flowcell.

### Bulk RNA-sequencing Analysis

All RNA-sequencing analyses were performed using integrated Differential Expression and Pathways Analysis, iDEP 2.01 (Ge et al., 2018). Genes expressed at extremely low levels were filtered out of the gene set by removing any genes with less than 0.4 counts per million (CPM) in at least one sample (n=1) (**Supplemental Figure 3A**). 13,934 filtered genes were converted to Ensembl gene IDs, normalized in edgeR and transformed via rlog. Hierarchical clustering of the samples and the top 2000 genes were performed (**Supplemental Figure 3B**). We then performed principal component analysis (PCA) using the 2000 most variable genes. Using DESeq2 and the Wald test (FDR cutoff = 0.1, minimum fold change = 1.5), we generated adjusted p-values and log_2_ fold changes. Gene expression was then compared between the “infected” AloCs and “control_uninfected” ALoCs. A total of 1,454 genes were found to be downregulated (log_2_ fold change <1) and 588 genes were upregulated (log_2_ fold change >1). Using analysis of variance (ANOVA) and post-hoc pairwise t-tests, we evaluated between-group differences for each cluster using these scaled expression values. We performed functional enrichment analysis using the hypergeometric test in hypeR for enriched pathways within the differentially expressed gene set. Enrichment analysis was run using significantly upregulated and significantly downregulated genes. Gene categories with fewer than 15 genes were excluded. Gene categories were considered significant if they had Benjamini-Hochberg adjusted *p*-values less than 0.1.

### Immunofluorescence

ALoCs were washed three times with 200 µL of 1X DPBS and 200 uL of 4% PFA in PBS were added to each channel of each ALoC for 20 min. at room temperature. ALoCs were then washed three times with 200 µL of 1X DPBS and incubated for 30 min. at room temperature with 200 µL of 0.1% Triton X-100 and 2% saponin. ALoCs were then washed three times with 200 µL of 1X DPBS and incubated overnight at 4°C with primary antibody (1:100) (**Supplemental Table 1**). ALoCs were washed again three times with 200 µL of DPBS and incubated with secondary antibody (1:1000) (**Supplemental Table 1**) for 2 hours at room temperature, protected from light. ALoCs were washed three times with 200 µL of 1X DPBS and incubated with DAPI for 10 min at room temperature. They were washed again three times with 200 µL of 1X DPBS and stored at 4°C with both channels filled with 1X DPBS. Confocal images were acquired on a Nikon CSU W1 spinning disk confocal microscope with an CFI S Plan Fluor ELWD 20x objective and a laser scanning confocal microscope Zeiss LSM 980 with an EC Plan-Neofluar 10x objective.

### Statistical analyses

All statistical analyses were performed using GraphPad Prism Software (version 9). For *in vitro* studies, data was analyzed using unpaired two-tailed *t*-test. For RNA sequencing analysis, analysis of variance (ANOVA) and post-hoc pairwise t-tests were used to evaluate between-group differences for each cluster using scaled expression values.

## Acknowledgements

The authors thank Gautam Mahajan and Ben Swenor from Emulate Inc. for providing protocols and support. The authors also thank core facilities at UT Southwestern Medical Center for their important contributions to this work including the UT Southwestern McDermott Center Next Generation Sequencing (NGS) Core and the Quantitative Light Microscopy Core, particularly the core director Marcel Mettlen, a Shared Resource of the Harold C. Simmons Cancer Center, supported in part by an NCI Cancer Center Support Grant, 1P30 CA142543-01 and Laura Alto of the UT Southwestern Microbiology Department Live Cell Imaging Facility, supported in part by an NIH S10 Instrumentation Award (1S10 OD034383).

## Data availability

Bulk RNA sequencing data has been deposited at the NCBI GEO database, GSE276053, and will be made publicly available at the time of publication.

## Author contributions

Conceptualization, V.E., M.U.S.; Formal analysis, V.E., M.U.S.; Funding acquisition, M.U.S.; Investigation, V.E., B.R., P.C.; Project administration, M.U.S.; Supervision, M.U.S.; Visualization, M.U.S.; Writing – original draft, V.E., M.U.S.; Writing – review and editing, All authors.

## Supplemental Figure Legends

**Supplemental Figure 1. Establishment of the Alveolus Lung-on-a-Chip Model**. **A.** Typical timeline of a single alveolus lung-on-a-chip (ALoC) experiment. **B.** Stitched image of the entire apical channel containing AT1 (HT1+, magenta) and AT2 cells (HT2+, green). Nuclei are labelled with DAPI (blue). Scale bar = 500 µm. **C.** A single z-slice image highlighting KRT5+ cells (orange) on an uninfected ALoC. Nuclei are labelled with DAPI (blue). Scale bar = 50 µm. **D.** A single z-slice image of the bottom of the basal compartment of an uninfected ALoC containing macrophages. Human lung microvascular endothelial cells (HMVEC) are labelled with VE-cadherin (magenta). Nuclei are labelled with DAPI. Scale bar = 50 µm. **E.** Stitched image of the entire apical channel of an ALoC containing macrophages (CD14+, orange), AT1 cells (HT1+, magenta), and AT2 cells (HT2+, green). Nuclei are labelled DAPI (blue). Scale bar = 500 µm. **F.** Brightfield image of an uninfected ALoC immediately after the addition human CD14^+^ peripheral blood-derived macrophages (macrophages) on top of the alveolar epithelial cells **(left)**. Brightfield image of an uninfected ALoC 24 hours after the addition of macrophages (**right)**.

**Supplemental Figure 2. Bulk RNA-sequencing analysis of *M. fortuitum*-infected ALoCs A.** Total raw read counts of uninfected (n=3) and *M. fortuitum*-infected (n=3) ALoCs. **B.** Pearson (average linkage) clustering of *M. fortuitum*-infected ALoCs and uninfected ALoCs.

