## Supplemental Figures for "An alveolus lung-on-a-chip model of *Mycobacterium fortuitum* lung infection"

**Supplemental Table 1.** Antibodies used on ALoC for immunofluorescence

| Cell Type | Target | Primary Antibody | Secondary Antibody |
|---|---|---|---|
| **AT1** | HT1 | Anti HT1-56 (Terrace Biotech, TB-29AHT1-56) | Alpaca anti-Mouse IgG1 Nano (VHH) Recombinant Secondary Antibody conjugated with Alexa Fluor 647 (Thermo Fisher, SA5-10333) |
| **AT2** | HT2 | Anti HT2-280 (Terrace Biotech, TB-27AHT2-280) | Goat anti-Mouse IgG2a Cross-Adsorbed Secondary Antibody, Alexa Fluor 488 (Thermo Fisher, A21131) |
| **Macrophage** | CD14 | CD14 Recombinant Rabbit Monoclonal Antibody (SC69-02) (Thermo Fisher, MA5-32248) | Donkey anti-Rabbit IgG (H+L), highly cross-adsorbed Secondary Antibody with Alexa Fluor 555 Plus |
| **Epithelial cell** | E-cadherin | E-cadherin recombinant rabbit monoclonal antibody (5H6L18) | Donkey anti-Rabbit IgG (H+L), highly cross-adsorbed Secondary Antibody with Alexa Fluor 555 Plus |
| **Endothelial cell** | VE-cadherin | CD144 (VE-cadherin) Monoclonal Antibody (16B1), eBioscience (Invitrogen 14-1449-82) | Alpaca anti-Mouse IgG1 Nano (VHH) Recombinant Secondary Antibody conjugated with Alexa Fluor 647 (Thermo Fisher, SA5-10333) |
| **n/a** | Nuclei | NucBlue™ Fixed Cell ReadyProbes™ Reagent (DAPI) (Thermo Fisher, R37606) | n/a |

### Supplemental Figure 1

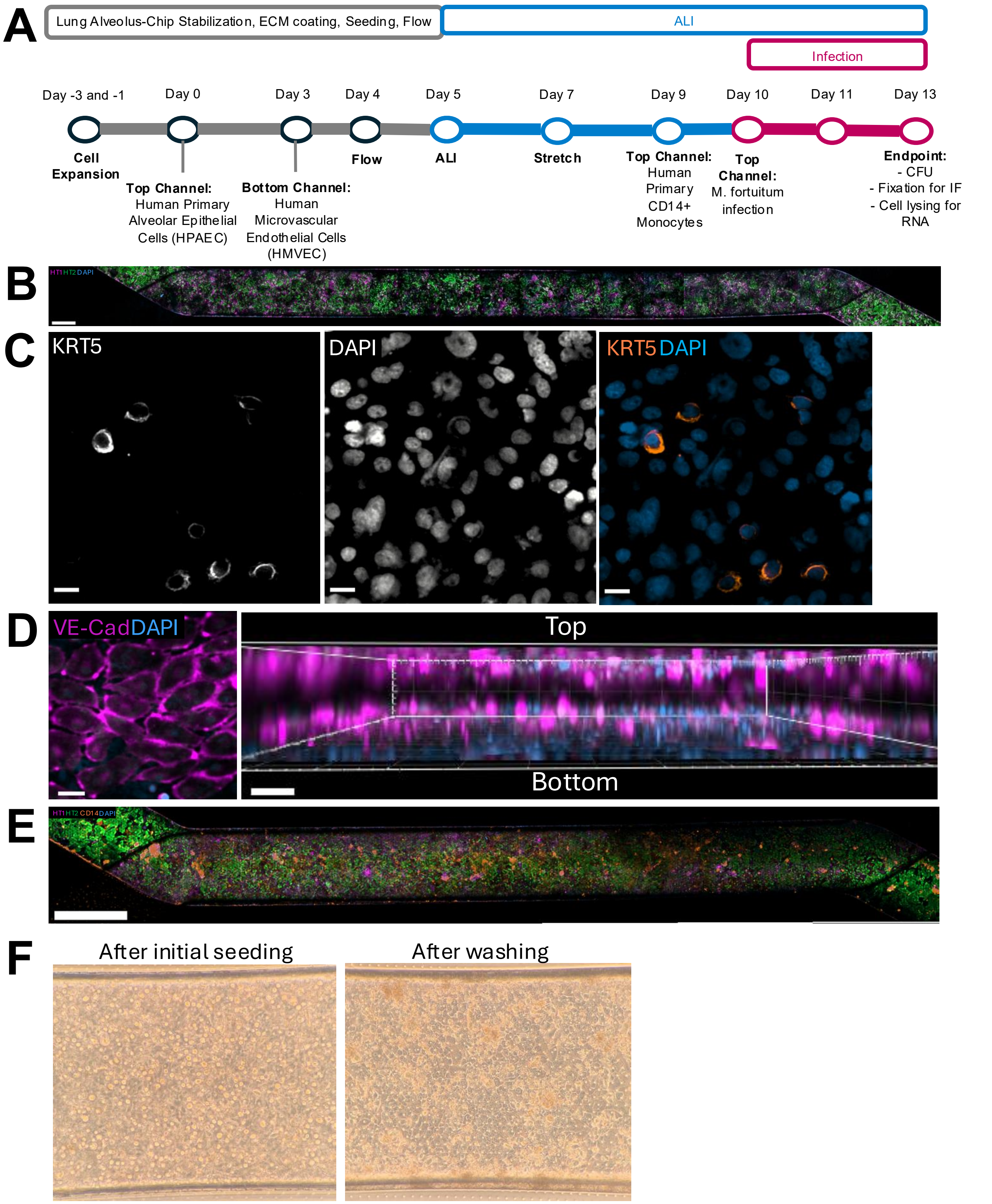

### Supplemental Figure 2

**A**

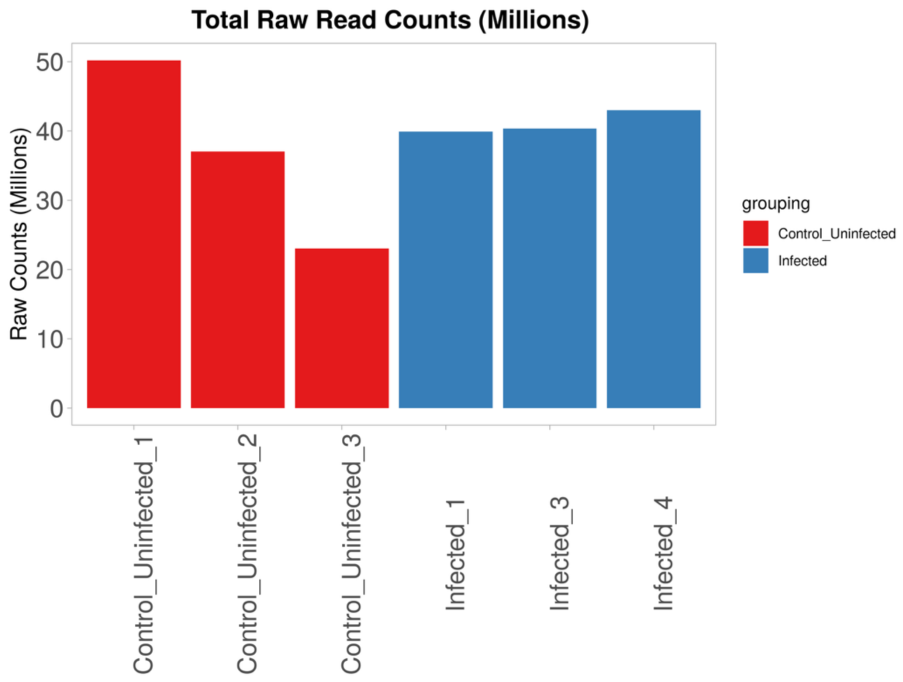


**B**

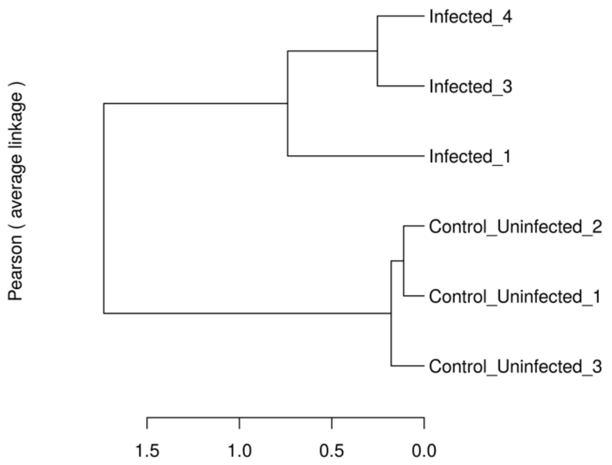
